# Probiotics enhance susceptibility of mice to cryptosporidiosis

**DOI:** 10.1101/304956

**Authors:** Bruno C. M. Oliveira, Giovanni Widmer

## Abstract

Cryptosporidiosis is a leading cause of diarrhea in infants and immune-compromised individuals. The lack of effective drugs against this enteric infection is motivating research to develop alternative treatments. To this aim, the impact of probiotics on the course of cryptosporidiosis was explored. The native intestinal microbiota of specific pathogen-free immunosuppressed mice was initially depleted with orally administered antibiotics. Then, a commercially available probiotic product intended for human consumption was added (or not) to the drinking water. Probiotic treated and untreated mice were orally infected with *Cryptosporidium parvum* oocysts. On average, mice treated with probiotic excreted more oocysts, indicative of a more severe infection. The probiotic treatment significantly altered the fecal microbiota, but taxonomic analyses showed no direct association between ingestion of probiotic bacteria and their abundance in fecal microbiota. These results suggest that probiotics indirectly alter the intestinal microenvironment in such a way that favors proliferation of *C. parvum*. The increase in the relative abundance of facultative anaerobes observed in mice with severe cryptosporidiosis indicates that dysbiosis is a consequence of severe cryptosporidiosis. The increase in the abundance of facultative anaerobes observed in severely infected animals is consistent with analyses of microbiota from individuals infected with other enteric pathogens. The results are significant because they show that *C. parvum* responds to changes in the intestinal microenvironment induced by a nutritional supplement.

**One Sentence Summary:** Mice treated with probiotics develop more severe symptoms of cryptosporidiosis.

## Introduction

Cryptosporidiosis is an enteric infection caused mostly by two species of *Cryptosporidium* parasites, *C. parvum* and *C. hominis*. Transmission occurs when infectious oocysts are ingested, either with contaminated food and water (1,2), by fecal-oral contact and possibly by inhalation (3). Recent surveys have revealed the high prevalence of cryptosporidiosis among infants living in developing nations, where it causes substantial morbidity and mortality in infants less than 2 years of age. (4). The treatment of cryptosporidiosis is limited to supportive care since no specific effective drugs available. Since no vaccines are available either, hygiene and water sanitation to reduce transmission remain the most effective approaches.

Using experimental mouse infections, we previous showed that cryptosporidiosis changes the gut microbiota. Given the importance of the microbiota to the physiology of the gut, here we investigated whether a reverse effect, of the microbiota on the parasite, could be demonstrated. We reasoned that the unmet need for anti-*Cryptosporidium* drugs could be alleviated by probiotics. This hypothesis does not necessarily imply that the microbial community of the gut directly impacts the parasite. Indeed, the transient nature of *Cryptosporidium* extracellular stages limits interactions between the resident microbiota and cryptosporidial life stages including sporozoites, merozoites and gametes. Alternatively, it is possible that the microbiota impacts parasite proliferation by modulating epithelial defense mechanisms, impacting the protective mucus layer, or stimulating innate and acquired immune cells.

The literature on the impact of the intestinal microbiota on cryptosporidiosis is sparse. A few studies have investigated the impact of *Cryptosporidium* parasites on the gut microbiota, but the effect of the gut environment on the course of the infection is not understood and the underlying mechanisms are unknown. Using germ-free immunodeficient SCID mice compared to SCID mice colonized with intestinal microbes, Harp et al. showed that normal intestinal microbiota delayed the onset of *C. parvum* oocyst excretion by several weeks (5). These authors also showed that resistance of mice to *C. parvum* can be increased by transferring intestinal mucosa from resistant animals to susceptible infant mice (6). A protective role of the gut microbiota against cryptosporidiosis was also observed in neonatal mice(7,8). These authors found that gut microbiota synergized with poly(I:C) to elicit a protective intestinal immunity against *C. parvum*. A study on the effect of inosine monophosphate dehydrogenase inhibitors in *C. parvum* infected mice detected an increase in *C. parvum* virulence in response to the drug. This effect was attributed to an alteration of the intestinal microbiota (9).

Research with probiotics in animal models of other infectious diseases has generated diverging results. A mouse model of rotavirus infection was used to show that administration of *Lactobacillus reuteri* reduced the duration of diarrhea (10). Similarly, and consistent with what has been observed in human trials, probiotics administered to mice had a mitigating impact on colitis induced by *Citrobacter rodentium* (11). The significant public health impact of nosocomial *Clostridium difficile* infection has generated a large body of research, including experiments in mice aimed at testing the benefit of fecal transplant (12) and defined probiotics (13–15). Only a few studies report a worse outcome with probiotic treatment. Using the cichlid fish tilapia, Liu et al. found that a 14-day treatment with probiotics made fishes more susceptible to infection with *Aeromonas hydrophila* after the treatment was discontinued (16). More relevant to the present study, research with mice showed that supplementation of diet with kefir exacerbated the outcome *Clostridium difficile* infection (17). Indicating that a harmful effect of probiotics is unusual, no other studies in rodent or mammalian models demonstrating increased susceptibility to infection appear to have been published.

We previously reported a significant impact of cryptosporidiosis on the profile of the bacterial intestinal microbiota (18). Replicated experiments with two *C. parvum* isolates comprising two infected and two control groups of mice revealed that the intestinal microbiota of infected animals differed from that of uninfected animals regardless of the *C. parvum* isolate. A taxonomic analysis of bacterial taxa highlighted two unclassified Bacteroidetes operation taxonomic unit (OTUs), Prevotellaceae and Porphyromonadaceae as overrepresented in the feces of infected mice, whereas OTUs most over-represented in uninfected mice were classified as Porphyromonadaceae and one unclassified Bacteroidetes OTU. With an eye on developing alternative treatments, the experiments described here were aimed at assessing whether probiotics can influence the course cryptosporidiosis. Although the results reveal a significant trend for more severe infection, the results show that *C. parvum* proliferation responds to changes in the intestinal microbiota. These observations open the way for targeted editing of the intestinal microbiota (19–21) as a low-cost approach to reducing the impact of cryptosporidiosis.

## Results

### Probiotic increases severity of infection

To test for a probiotic effect on the severity of *C. parvum* infection, fecal oocyst output was measured by flow cytometry as described above. In experiment 1, a total of 92 oocyst concentration values were acquired from 16 mice and six timepoints over a 15-day period. In experiment 2, 79 datapoints were obtained from the same number of mice (Fig. 1). In experiment 1 mice which received probiotic excreted a significantly higher concentration of oocysts. A mean oocyst output of 76,463 oocysts/g feces (n =44) was measured against a mean of 26,732 oocysts/g for the control mice (n=48). The difference between treatments was highly significant (Mann-Whitney U=541, p<0.001). An analogous significant probiotic effect was obtained for experiment 2; mean_probiotic_=378,736 oocysts/g, n_probiotic_=37, mean_control_=68,778 oocysts/g, n_control_=42; U=269.5, p<0.001). Fig. 1 shows the pattern of oocyst output for the two experiments plotted on a log scale. In experiment 1, due to early mortality and high oocyst output, the concentration of dexamethasone in the water was lowered from 16 mg/l to 10 mg/l on day 8.

**Fig. 1.**
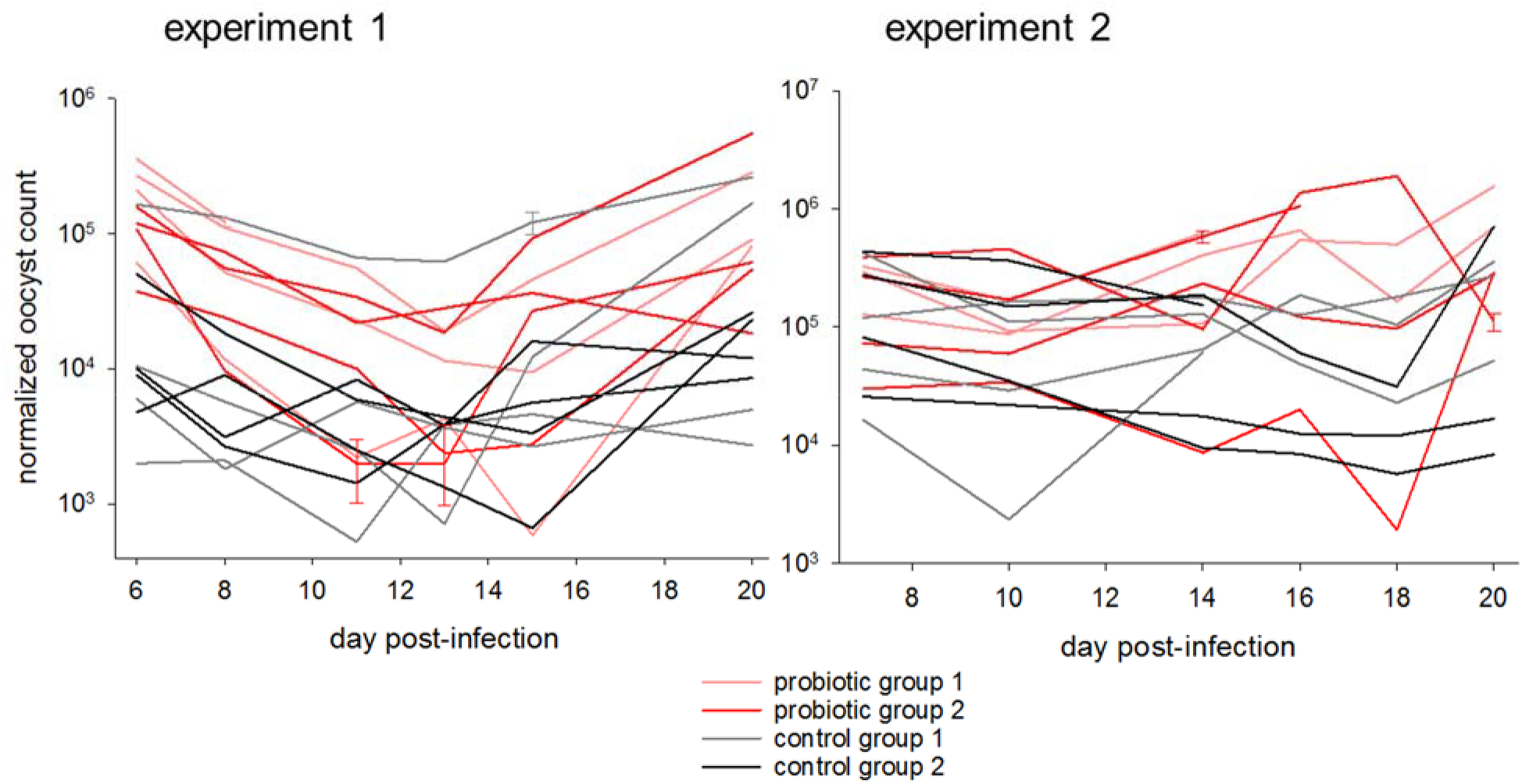
Effect of probiotic on severity of *C. parvum* cryptosporidiosis in immune-suppressed mice. The graphs show normalized oocyst counts expressed as oocysts/g feces for two independent experiments. Each line represents one mouse. Error bars show SD based on five replicate counts of selected samples.

This intervention is the likely cause of the reduction in oocyst output. Mouse mortality appeared to be unrelated to probiotic treatment, but the numbers are too small to support statistical testing. In experiment 1, two mice out of 16 died during the experiment; one mouse from treatment group 1 on day 8 PI, and one mouse from control group 1 on day 15 PI. In experiment 2 two mice died, from treatment group 2 on day 16 PI and one from control group 1 on day 14 PI.

### Probiotic treatment significantly impacts fecal microbiota

To assess whether probiotic treatment impacted the fecal microbiota, pairwise weighted UniFrac distances (22) between sequence data from 64 experiment 1 fecal samples were visualized on a PCoA plot (Fig. 2). Samples collected starting on day 5 of probiotic treatment (day 4 PI) until day 16 of treatment (day 15 PI) were included. The number of samples in this analysis is smaller than shown in Fig. 1, because not all fecal samples were sequenced. Consistent with an impact of probiotic consumption on the profile of the intestinal microbiota, this analysis revealed a nonoverlapping distribution of datapoints according to treatment. ANOSIM was used to check the significance of sub-structuring by treatment. The test returned a highly significant *R* value of 0.305 (p<10^−5^). Analogous results were obtained for experiment 2 based on 55 samples collected starting on day 10 after probiotic treatment was initiated (day 9 PI) until day 20 (day 19 PI). As for experiment 1, clustering by treatment was significant (R = 0.210, p<10^−5^).

**Fig. 2.**
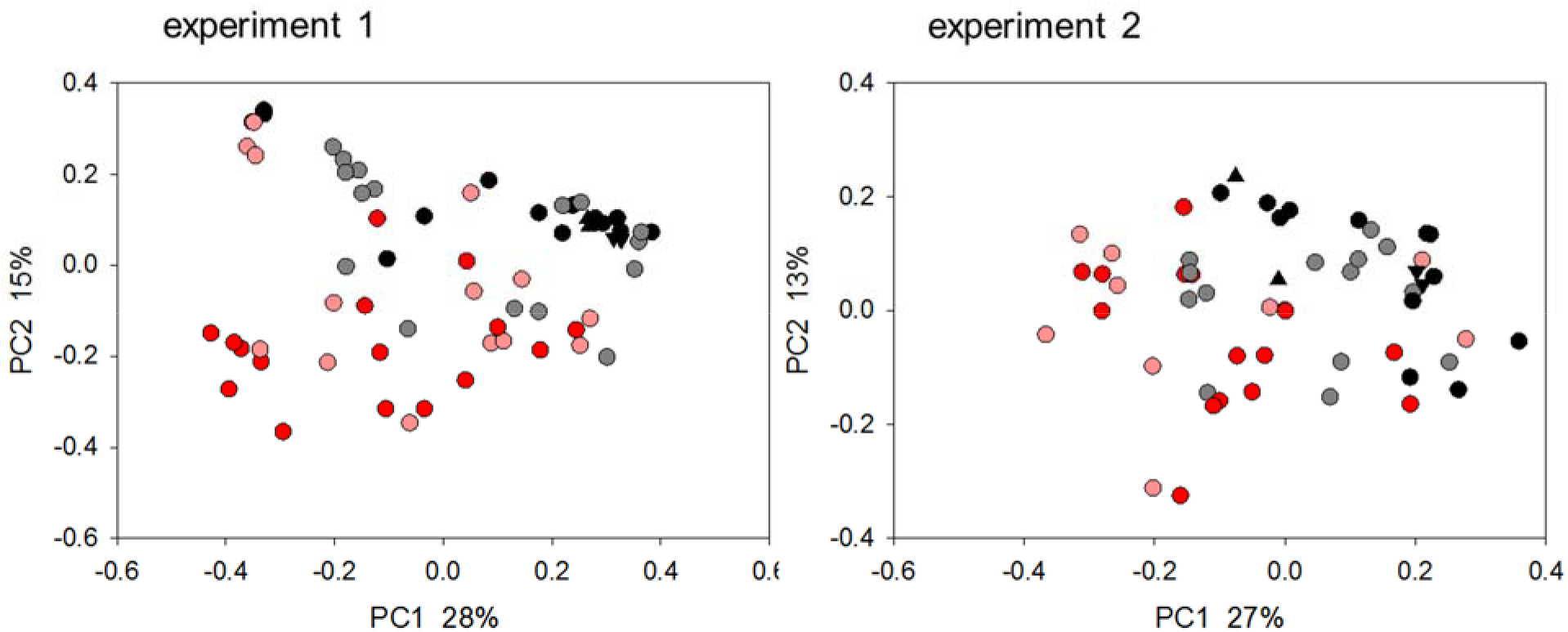
Impact of probiotic on the fecal microbiome of *C. parvum* infected mice. Principal Coordinate Analysis was used to display weighted UniFrac distances between pairs of fecal microbiome samples. Experiment 1 analysis (left) includes data from 64 fecal samples collected from day 5 of treatment (day 4 PI) until day 16 of treatment (day 15 PI). For experiment 2, 55 samples were analyzed. Each datapoint represents one sample, color-coded according to treatment and group as shown in Fig. 1. Matching triangle symbols indicate replicate analyses of the same fecal samples.

### Bacterial α-diversity does not correlate with oocyst output

Having detected an impact of probiotic consumption on oocyst output and on the fecal microbiota, the FCM and 16S data were analyzed jointly to identify possible associations between bacterial microbiota profile and severity of cryptosporidiosis. These combined analyses included all samples for which 16S and FCM data were acquired. A total of 44 samples were included in each experiment. As dysbiosis is typically characterized by low bacterial diversity and is often associated with increased susceptibility to enteric infections (23) the first global analysis examined 16S sequence data and oocyst concentration for a possible correlation between microbiota α-diversity and total oocyst output (Fig. 3). Regardless whether all data were pooled by experiment, or samples from probiotic treated and control mice were analyzed separately, the correlation was very low, explaining 9% of oocyst output at most. If considering only the 10 samples with the highest and lowest oocyst concentration (20 samples per experiment), no correlation between oocyst output and α-diversity was apparent either. We also used the Mantel test to assess whether the difference in oocyst output between pairs of samples correlated with UniFrac distance. In other words, with this analysis we tested whether samples harboring very different concentrations of oocysts also exhibited large phylogenetic distances. No association between these variables was identified in either experiment (R_xy_=0.12, n=44, p=0.055 and Rxy=0.058, n=53, p=0.28). Together with the results shown in Fig. 3, these analyses indicate that higher oocyst output is not correlated with a global change in the microbiota.

**Fig. 3.**
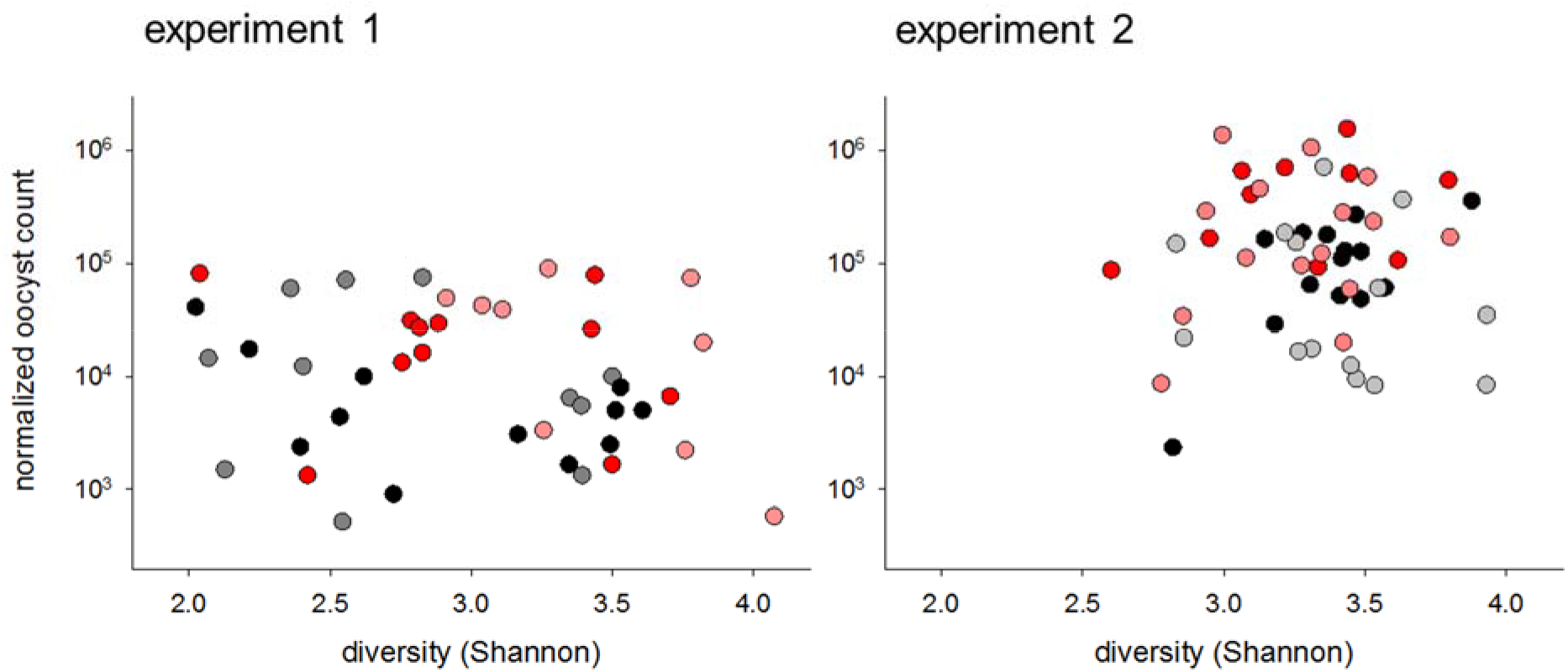
Oocyst output is unrelated to diversity of bacterial microbiome. Oocyst output normalized for feces volume was plotted against Shannon diversity for 44 samples collected on three days which were analyzed for both properties in experiment 1 and 53 samples from experiment 2. Color indicates experimental group as described in Fig. 1. Samples were collected from individual mice.

RDA was used as the second global analysis to assess whether fecal oocyst concentration significantly correlated with microbiota profile and identifying bacterial OTUs correlating in relative abundance with oocyst output. In experiment 1 a Monte Carlo test involving 1000 permutations of the 44 samples indicated a significant correlation between oocyst concentration and OTU profile (pseudo-F=1.0, p=0.014). This effect remains significant (F=2.7, p=0.024) after accounting for treatment, i.e., defining treatment (±probiotic) as covariate, or accounting for “mouse” (defining mouse as covariate; pseudo-F=3.8, p=6×10^−5^). The analogous test in experiment 2 also returned a significant pseudo-F ratio of 4.2 (n=44, p=0.0016). If removing the effect of treatment or “mouse” by defining each variable as covariate, the association remains significant (pseudo-F=2.6; p=0.015 and pseudo-F=5.2; p=3×10^−4^, respectively).

### High abundance of facultative anaerobes in severe infections

Having identified a significant correlation between fecal oocyst concentration and OTU profile, the taxonomic make-up of the fecal microbiota was examined in more detail with the goal of identifying bacterial taxa which correlate in abundance with oocyst output. First we used program LEfSe (24) to identify OTUs which significantly define the difference between samples containing high and low oocyst concentration. This analysis was based on 20 samples for each experiment, 10 samples with the highest oocyst concentration and 10 samples with the lowest concentration. In experiment 10, 7/10 samples in the high-oocyst group originated from mice treated with probiotics and 7/10 samples in the low-oocyst group came from control mice (Chi-square=3.2, p=0.07). In experiment 11, 9/10 samples in the high-oocyst group originated from treated mice and 8/10 samples in the low-oocyst group originated from control mice (Chi-square=9.9, p=0.002). Feces from highly infected animals were characterized by a high Proteobacteria abundance, whereas feces from lightly infected animals were significantly enriched for Firmicutes. As observed for the impact of the probiotic on the severity of cryptosporidiosis (Fig. 1) and on the global microbiota profile (Fig. 2), a clear similarity was observed between the two experiments (Fig. 4).

**Fig. 4.**
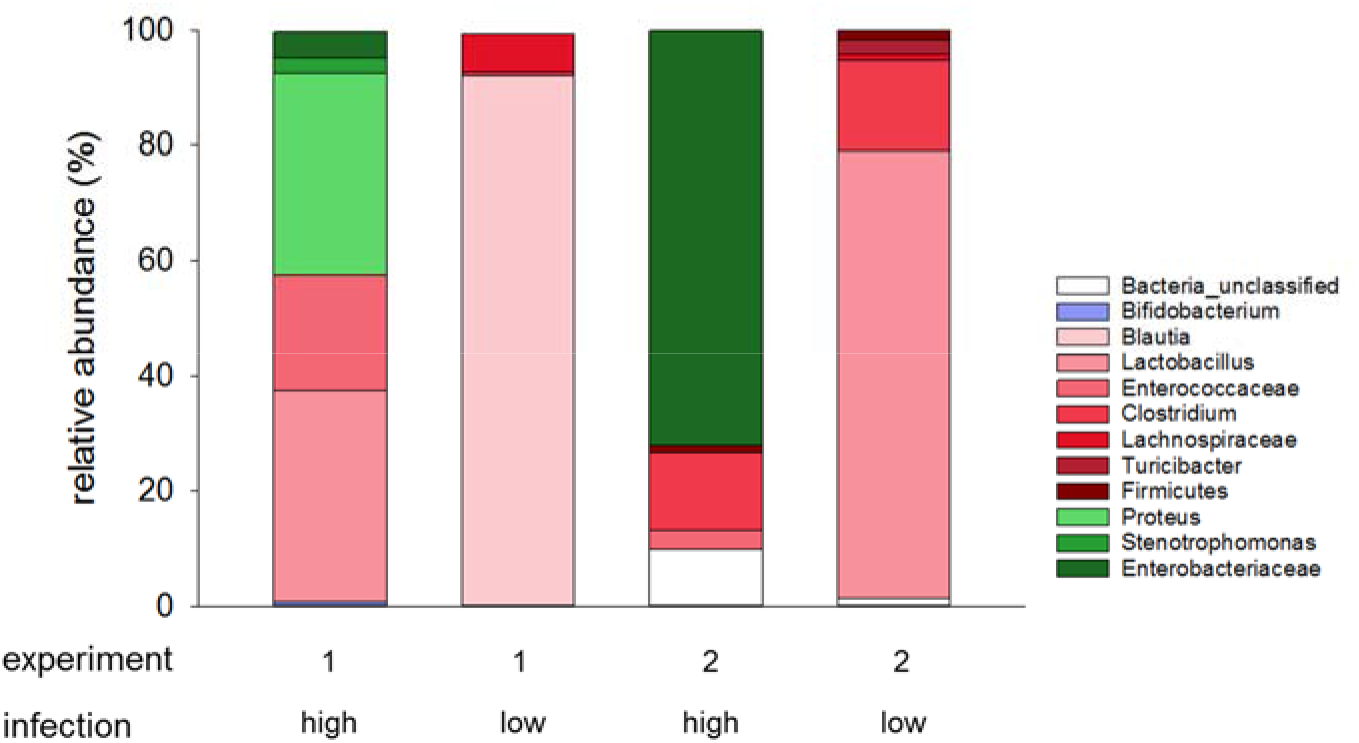
Taxonomy of fecal microbiome from heavily and lightly infected mice shows that heavy infections are associated with increased abundance of Proteobacteria. For each experiment, bacterial taxa significantly associated with severity of infection were identified using LEfSe (24) in a comparison of ten samples with the highest (high) and lowest (low) oocyst concentration; twenty samples were included for each experiment. Color indicates phylum and color intensity genus or highest taxonomic level (family, order etc.) identified; green Proteobacteria; red, Firmicutes; blue, Actinobacteria.

A second taxonomy analysis aimed at identifying OTUs enriched in highly infected animals was performed with RDA (25). For experiment 1, in the 10 OTUs which best correlated in relative abundance with oocyst concentration, 61% of the sequences were classified as Lactobacillus, 24% as Proteus and 14% as Enterococcus. In the 10 OTUs which most negatively correlated with oocyst concentration, as seen in with LEfSe, *Blautia* was the most abundant classification (61% of reads), followed by Clostridiaceae (27%), Lachnospiraceae (10%), Romboutsia (8%) and unclassified Firmicutes. For experiment 2, the identical analysis of the 10 OTUs which best correlated in relative abundance with oocyst concentration, 92% of sequences originated from Enterobacteriaceae and 8% from Firmicutes. In the 10 OTUs with lowest oocyst concentration 70% of sequences were classified as Lactobacillus, 21% as Turicibacter, 6% as unclassified Lactobacillales and 1% as Stenotrophomonas (phylum Proteobacteria). The results obtained with RDA are thus is close agreement with the LEfSe results shown in Fig. 4.

### Severity of infection correlates with fecal microbiota profile across experiments

Fig. 3 illustrates the main difference the experiments; mean fecal oocyst concentration across all time points, treatments and mice in experiment 2 was 2.4 × 10^5^ oocysts/g (n=79, SD=3.4 × 10^5^), or about 5 times higher than in experiment 1 (mean=5.0 × 10^4^ oocysts/g, n=92, SD=8.7 × 10^4^; Mann-Whitney U=1467, p<0.001). It is likely that this difference was caused by the different routes of dexamethasone administration: drinking water only in experiment 1, vs. drinking water followed by IP injection in experiment 2. The more severe infection in experiment 2 represents an unplanned opportunity to further assess the impact of the infection on the gut environment. If heavier infections cause an increase in Proteobacteria relative abundance, as suggested by LEfSe analysis (Fig. 4) and RDA described above, one would expect a higher proportion of Proteobacteria in experiment 2 samples originating from severely infected mice, which is exactly what was observed (Fig. 4 and RDA results). Further, severe cryptosporidiosis can be expected, based on research on human patients suffering from other enteric infections (26–28), to lead to a change in bacterial microbiota towards populations enriched for facultative anaerobes, possibly resulting in less diversity. To test this hypothesis, mean pairwise weighted UniFrac distances between microbiota from heavily infected samples were compared to distances between lightly infected samples. In experiment 1, mean β-diversity between the 10 high-oocyst samples was 0.536 (45 pairwise distances, SD=0.142), whereas between low-output samples mean β-diversity was 0.590 (45 pairwise distances, SD=0.121; Mann-Whitney T=1819.0, p=0.07). In experiment 2, the mean β-diversity values are 0.312 (45 pairwise distances, SD=0.117) and 0.411 (45 pairwise distances, SD=0.137) for the 10 samples with the highest and lowest oocysts concentration, respectively (Mann-Whitney U=590, p<0.001). Although for experiment 1 the effect is not significant, together these results are consistent with the model postulated above; i.e. that severe infection leads to a convergence of the microbiota, towards an increase in the relative abundance of Proteobacteria and a reduction in Firmicutes.

### Loss of microbiota functional diversity in heavily infected mice

To extend the observed taxonomic differences between severely and lightly infected mice to the metagenome, program PICRUSt (29) was used to infer microbiota function from OTU profiles. Given the more severe infection in experiment 2, metagenome analyses are only reported for this experiment. PICRUSt identified 39 KEGG level 2 categories in the combined metagenome. A PCoA based on pairwise SSR distance between KEGG abundance values normalized across KEGG categories revealed a tight clustering of samples with high oocyst concentration relative to the samples with lower oocyst concentration (Fig. S1). This visual assessment was tested by comparing pairwise distances between KEGG profiles. For the ten samples with the lowest oocyst concentration the distance averaged 69.9 (SD=47.6) and for the same number of samples with the highest concentration 54.8 (SD=41.41), which is statistically not significant (MannWhitney Rank Sum test, p=0.121). If only eight samples with highest and lowest oocyst concentration were tested (28 pairwise distance values for each group), the distances between the high concentration samples are significantly smaller (Mann-Whitney Rank Sum test p=0.015).As for the taxonomy analysis described above, we conclude from these results that proliferation of *C. parvum* leads to a convergence of the inferred bacterial metagenome.

As described above for the taxonomy analysis, LEfSe was used to identify KEGG pathways which differ significantly in abundance between the 10 samples from highly infected and the same number of samples from lightly infected mice. Underscoring the difference at the metagenome level between fecal samples from severely and mildly infected mice, 22 of 39 KEGG level 2 pathways were significantly different between the two groups (Table S1). In comparison to the severely infected mice, the microbiota of mild infections was characterized by a high abundance of pathways related to replication, such as carbohydrate, amino acid and nucleic acid metabolism. These results extend the taxonomy presented in Fig. 4, suggesting that the mouse dysbiotic cryptosporidiosis metagenome is selected for other functions than bacterial replication.

## Discussion

In a comprehensive review of the literature, Kristensen et al. (30) found a small number of publications describing randomized controlled probiotics trials which included the characterization of fecal microbiota. The surprising conclusion of this survey is that no publications reported a significant change in the microbiota based on OTU richness, evenness or diversity analysis. Although our experiments were not designed to assess the probiotics’ impact on the microbiota, we did find a significant probiotic effect. However, as no uninfected groups treated with probiotics were included, the change in microbiota profile could be related to the severity of cryptosporidiosis rather than a direct impact of consumption of probiotics. Our focus was not to model the impact of probiotics on the gut ecosystem, but to assess whether probiotics can mitigate the severity of cryptosporidiosis.

As we do not observe a significant increase in probiotic bacteria in the feces of treated mice, we postulate that some of the bacterial or prebiotic ingredients present in the probiotic product induced changes in the mouse intestinal environment favoring the proliferation of *C. parvum*. Proliferation of the parasite then leads to extensive secondary modifications of the microbiota which were detected by 16S amplicon sequencing. An impact of the prebiotics present in the product, acacia gum, larch gum, oligosaccharides and L-glutamine, on the microbiota cannot be excluded. Elucidating the mechanism by which probiotic administration promotes proliferation of *C. parvum* will require testing of individual probiotic species or defined combinations of species and/or prebiotics (31) and metabolomics analysis to identify potential mediators of the probiotics effect. This research is of primary importance to enable targeted manipulations of the microbiota aimed at limiting proliferation of *Cryptosporidium* parasites. Zhu et al. (19) describe methods to “edit” the gut microbiota, in this case by inhibiting the multiplication of facultative anaerobes. An analogous approach could be used to investigate the causal link between parasite proliferation and dysbiosis. Although mice infected with *C. parvum* do not develop diarrhea, the fecal microbiota from heavily infected animals in our experiments resembles the fecal microbiota of humans suffering from cholera diarrhea (26–28) or diarrhea of other etiologies (28), indicating that neither pathogen type nor fecal water content are important to change the microbiota. A characteristic of many intestinal pathologies of infectious or other causes is an increase in the proportion of Gammaproteobacteria (32). Although exceptions to this trend have been reported (33), a shift towards facultative anaerobes reflecting increased permeability of the gut epithelium is a hallmark of infectious (26,27,34,35), inflammatory (36–38) and other intestinal pathologies (39). The abundance of Gammaproteobacteria in the distal gut of mice heavily infected with *C. parvum* is significant because it indicates a shift in the luminal oxygen gradient (40) creating conditions favoring multiplication of Gammaproteobacteria. Thus, even in the absence of diarrhea, the mouse is a valuable model of cryptosporidiosis pathology. The observed abundance of facultative anaerobes in the absence of diarrhea also demonstrates that diarrhea is not a direct cause of dysbiosis. Previous observations showing that cryptosporidiosis impacts the murine microbiota in the absence of any specific treatment (18) is consistent with dysbiosis being a consequence of the infection, perhaps caused by epithelium erosion, villus atrophy (41–43), increased sloughing of epithelial cells and a resulting increase in luminal O_2_. These observations raise the question whether *Cryptosporidium* proliferation responds to oxygen concentration in the gut lumen. Although the proliferative stages of the parasite are intracellular, sporozoites, merozoites and microgametes are extracellular. Selective inhibition or promotion of oxygen-consuming bacteria (19) to temporarily raise or deplete luminal O_2_ (20) could potentially be investigated to assess the response of *C. parvum* and explore indirect approaches to mitigating the severity and duration of cryptosporidiosis through microbiota manipulation.

Few studies have reported on the effect of diet on *C. parvum* infection. Liu et al. (44) found that protein deficiency increases the concentration of *C. parvum* DNA in feces. However, the difference between normal and protein-deficient animals was reported at 20 h post-infection. Since *C. parvum* is not known to complete its life cycle in less than 72 h (45), the diet effect is difficult to interpret. A second study found a positive effect of pomegranate extract on cryptosporidiosis in calves (46). The authors report that calves fed milk supplemented with extract excreted fewer oocysts. The very limited range of the literature on the effects of diet on cryptosporidiosis illustrates the need for additional research, particularly basic research on mechanistic aspects of parasite-microbiota interaction. The possibility of including the innate and adaptive immune response in this research is currently limited by the fact that studies in mice infected with *C. parvum* require immune suppression.

Although the results of experiment 1 and experiment 2 are consistent, differences were also noticed. Most notably, average oocyst output in experiment 2 was higher, indicative of a more severe infection. Corroborating the model discussed above, more severe infections were associated with higher relative abundance of Gammaproteobacteria (Fig. 4). The reason for the difference between severity of infection is hard to determine, but the different route of dexamethasone administration mentioned above could have contributed to this outcome. The use of two *C. parvum* isolates is unlikely to be a relevant factor as MD and TU114 are not known to differ with respect to their virulence phenotype in dexamethasone treated mice (unpublished observations). Differences in fecal oocyst output between co-housed mice was also observed (Fig. 1). As described in Materials and Methods, animals from a same cage were housed individually only for 16 h three times a week for collection of feces, but were otherwise housed together in two cages per treatment or 4 cages for each experiment. The difference in oocyst output and microbiota profile between cagemates is difficult to explain given the close contact between animals. This phenomenon justifies mice to be sampled individually, rather than sampling by group as is common practice.

## Conclusions

In the absence of effective anti-protozoal drugs to control cryptosporidiosis, alternative treatments are attractive. However, our current understanding of how microbiota perturbation, whether induced by diet, pro-, anti- or prebiotics, affects enteric pathogens including *Cryptosporidium* is very incomplete. Identifying specific mechanisms impacting pathogen virulence in response to probiotics consumption or diet may enable the development of targeted microbiota editing measures to mitigate the severity of cryptosporidiosis. Methods designed to detect metabolome modifications (47) will be needed to supplement the information gained from 16S amplicon sequencing. Lastly, enhancing the value of the rodent cryptosporidiosis model, the observed shift towards facultative anaerobes indicates common pathogenic changes in the human and rodent intestine in response to enteric infections.

## Materials and Methods

### Parasites

Oocysts from *C. parvum* isolate MD (48) or TU114 (49) were used in experiment 1 and 2, respectively. MD is a zoonotic isolate, whereas TU114 belong to the anthroponotic subgroup characterized by a IIc GP60 genotype (50). Oocysts were purified from mouse feces on gradients of Nycodenz (Alere Technologies, Oslo, Norway) as described (51). The age of the oocysts was 13 and 110 days for experiment 1 and 2, respectively.

### Mouse experiments

To test the effect of a commercially available probiotic, two replicate experiments were conducted in mice. In each experiment, herein referred to as experiment 1 and experiment 2, respectively, 16 female CD-1 mice approximately 6 weeks of age were used. Upon delivery, each mouse was individually tagged and randomly assigned to one of four groups of four mice. Mice were immunosuppressed by adding dexamethasone 21-phosphate disodium (Sigma, cat no. D1169) to the drinking water at a concentration of 16 mg/l (52) starting on the day of arrival, defined here as day −7 post-infection (PI), i.e., 7 days prior to infection. To deplete the native intestinal microbiota, vancomycin and streptomycin were added to the drinking water at a concentration of 500 mg/l and 5 g/l, respectively, starting on day −6 PI. Metronidazole at a dose of 20 mg/kg was given daily by gavage starting on day −6 PI. The antibiotic treatment was discontinued on day −2 PI. Starting on day −1 PI, the drinking water was supplemented with 1.3 g of probiotic added to 500 ml water. This product contains 15 bacterial strains belonging to the genera *Bifidobacter* (4 species), *Lactobacillus* (9 species) and *Streptococcus thermophilus*. In addition to bacteria, 1 g of product contains 8 mg Acacia gum, 540 mg Larch gum, 115 mg galactooligosaccharide, 212 mg L-glutamine and 150 IU vitamin D3. When dissolved into 500 ml of water, the final concentration of these ingredients was 20 μg/ml, 1.4 mg/ml, 0.3 mg/ml, 0. 55 mg/ml and 0.4 IU/ml, respectively. The product is flavorless. Drinking water with dexamethasone and probiotic water was replaced every 3 days. Mice were orally infected on day 0 PI with approximately 5 × 10^4^ *C. parvum* oocysts. In experiment 1, to compensate for increased water uptake in the two groups receiving probiotic, the concentration of dexamethasone phosphate in the water of the two treated groups was reduced to 10 mg/l starting on day 10 PI. In experiment 2, to avoid possible differences in dexamethasone uptake with drinking water, the drug (Dexamethasone; Sigma, cat no. D1756) was given only subcutaneously every second day starting on day 2 PI. A volume of 100 μl of 10 mg/ml dexamethasone suspension was injected alternatively into the right and left side of the abdomen. To obtain fecal pellets for microbiota analysis, mice were individually transferred to a 1-liter plastic beaker and pellets collected upon defecation. Pellets were stored at −20°C. To collect feces for oocyst enumeration, mice were individually transferred overnight to collection cages fitted with a wire bottom. Feces were collected from these cages were stored at 4°C. In the morning, mice were returned to their respective group cage, such that they were housed individually for 14-16 h on the days when feces for oocyst enumeration were collected. While in conventional cages, the mice were always housed with the same cagemates.

Animal experiments adhered to the NIH Guide for the Care and Use of Laboratory Animals and were approved by the Tufts University Animal Care and Use Committee.

### Enumeration of oocysts

Feces collected overnight were weighted, diluted 1:5 in distilled water and homogenized with a vortex. To remove debris, fecal homogenates were filtered through 100 μm cell strainers (Corning cat. no. 431752) by centrifugation at 1300 × g. The filtrates were homogenized and volumes of 1 ml centrifuged at 6700 × g for 5 min. The supernatant was discarded and the pellet was suspended in 500 μl of PBS supplemented with 10% fetal bovine serum. A volume of 20 μl of this suspension was transferred to a 1.5 ml microcentrifuge tube and 20 μl of a 1:5 dilution of monoclonal antibody 5F10 cell culture supernatant. This antibody reacts specifically with the *Cryptosporidium* oocyst wall without binding to other parasite antigens (Sheoran, unpublished observation). Samples were incubated for 30 min at room temperature. Following incubation with primary antibody, the samples were centrifuged for 10 min at 6700 × g, the pellet washed once in 500 μl of PBS with 10% FBS and incubated with 20 μl of secondary antibody (Alexa Fluor 488 goat IgG anti-mouse) for 30 min at room temperature. After incubation, 500 μl of PBS was added and the samples washed once in PBS. For each experiment five samples were randomly selected for replication. Replication involved processing and labelling 5 separate aliquots originating from each strained and washed sample. Labelled samples were analyzed by flow cytometry using a Becton Dickinson Accuri C6 cytometer. Distance matrices were calculated in GenAlEx 6.5 (53) based on the pairwise difference between oocyst concentrations. Specifically, the distance between sample A containing an oocyst concentration of x_a_/g feces and sample B containing a concentration of x_b_/g feces was calculated as (x_a_ - x_b_)^2^. The accuracy of the flow cytometry oocyst enumeration method was evaluated by correlating the counts against the *Cryptosporidium* 18S ribosomal RNA gene copy number estimated using real-time PCR with published primers (54).This analysis based on five randomly chosen experiment 1 samples generated a correlation coefficient of 69%.

### Microbiota analysis

DNA was extracted from 10 mg of feces collected individually from each mouse. DNA was extracted in a QIACube instrument using the QIAamp PowerFecal DNA kit (QIAGEN, cat. 12830-50) according to manufacturer’s protocol. DNA was eluted in 50 μl of elution buffer and stored at −20°C. A previously described PCR protocol to prepare 16S V1V2 amplicons libraries for high-throughput sequencing was used (18). The only deviation from this procedure was the downstream primer; instead of canonical primer 338R we used primer Bac R V2 short (GTTCAGACGTGTGCTCTTCCGATCtgctgcctcccgtaggagt), where the lowercase characters are equivalent to the conserved 338R sequence (55). One microliter of primary PCR reaction was subjected to a secondary PCR to incorporate a 6-nucleotide barcode using to each sample. The secondary amplification was as described (18), except that downstream primer *CAAGCAGAAGACGGCATACGAGAT*nnnnnnGTGACTGGAGTTCAGACGTGTGCTCTTCC was used. The italicized nucleotides represent the Illumina adaptor and the lowercase characters the barcode unique to each sample. To assess the quality of the PCRs, a portion of the amplification product was electrophoresed on 1.5% agarose and visualized using GelRed™ (Biotium). The concentration of each final amplicon was measured in a Qbit spectrophotomer and up to 80 amplicons pooled at approximately equal concentration. The pooled library was size selected with a Pippin HT library size selection system (Sage BioScience). Libraries were sequenced in an Illumina MiSeq sequencer at Tufts University Genomics core (tucf.org) using single-end 300-nucleotide strategy. To control for technical variation introduced during library preparation and sequencing, each library included two replicates of two randomly chosen samples. Replication involved the processing of duplicate fecal samples processed, amplified and barcoded individually.

### Bioinformatics

FASTQ formatted sequences were processed using programs found in mothur (56) essentially as described (18). Briefly, random subsamples of 5000 sequences per sample were processed. This procedure is not expected to bias the analysis (57) since the number of sequences per barcode was relatively constant. The mean number of sequences per sample in experiment 1 was 1.03 × 10^5^ (SD=2.44 × 10^4^). In experiment 2 the average number of reads was 1.01 × 10^5^ (SD=3.22 × 10^4^), representing a coefficient of variation of 0.23 and 0.31, respectively. V1V2 sequences were trimmed to 200 nucleotides to eliminate 3’ sequence with a mean Phred quality score <30. Sequences were aligned and sequences with the following properties were removed: sequences that did not align, sequences with ambiguous base calls and sequences with homopolymers >8 nt. To further reduce the number of putative sequence errors, program *pre.cluster* was used to merge unique sequences differing by one nucleotide position with the majority sequence (58). This sequence curation protocol removed 15,748 from 400,000 experiment 1 sequences (3.9%) and 21,546 from 355,000 experiment 2 sequences (6.0%) sequences. Pairwise UniFrac phylogenetic distances (22) between samples were calculated in *mothur*. Analysis of Similarity (ANOSIM) (59) was used to test the significance of clustering by treatment. Program *anosim* was run in *mothur* using a weighted UniFrac distance matrix as input. Operational Taxonomic Units (OTUs) were obtained using program *cluster*, also found in *mothur*, using the OptiClust clustering method (60) and a distance cut-off of 3%.

Linear Discriminant Analysis as implemented in program LEfSe (24) was used to identify statistically significant differences in OTU abundance profiles between two groups of samples defined indepently of treatment as heavily and lightly infected. Heavy and light infection was defined on the basis of fecal oocyst concentration determined by flow cytometry as described above. In the experiment-wide LefSe analyses, the 10 samples with the highest oocyst concentration and the 10 samples with the lowest oocyst concentration were selected. When analyzing subsamples of probiotic and control mice separately, the five samples with the highest oocyst concentration and the same number of samples with the lowest oocyst concentration were tested using LEfSe (24).

Redundancy Analysis (RDA) was used to test the significance of association between OTU profile and oocyst concentration or between KEGG function profile and oocyst concentration. The program was run in CANOCO (25). OTU abundance values for the 100 most abundant OTUs, or KEGG functions (n=39) inferred with metagenome prediction tool PICRUSt (29) served as dependent variables. Oocyst concentration determined by flow cytometry as described above served as the independent variable. Pairwise distance between KEGG profiles was calculated as described above for oocyst concentration, except that the squared difference was summed over all KEGG categories. This metric is equivalent to the square of the Euclidean distance.

Correlation between pairs of matrices was tested with the Mantel test (61). This test uses random permutation of the elements in one matrix to detect a relationship between the elements of two matrices (62). The implementation of Mantel in GenAlEx 6.5 (53) was used.

16S sequence data from experiment 1 and experiment 2 were deposited in the ENA Sequence Read Archive under study accession number PRJEB25162 and PRJEB25164, respectively.

## Acknowledgements

Lucas Vinicius Shigaki de Matos, Olga Douvropoulou and Kevin Huynh performed preliminary experiments.

## Author contributions

BCMO and GW designed the experiments and analyzed the data. BCMO performed the experiments. GW wrote the manuscript.

## Supplementary materials

**Fig. S1.**
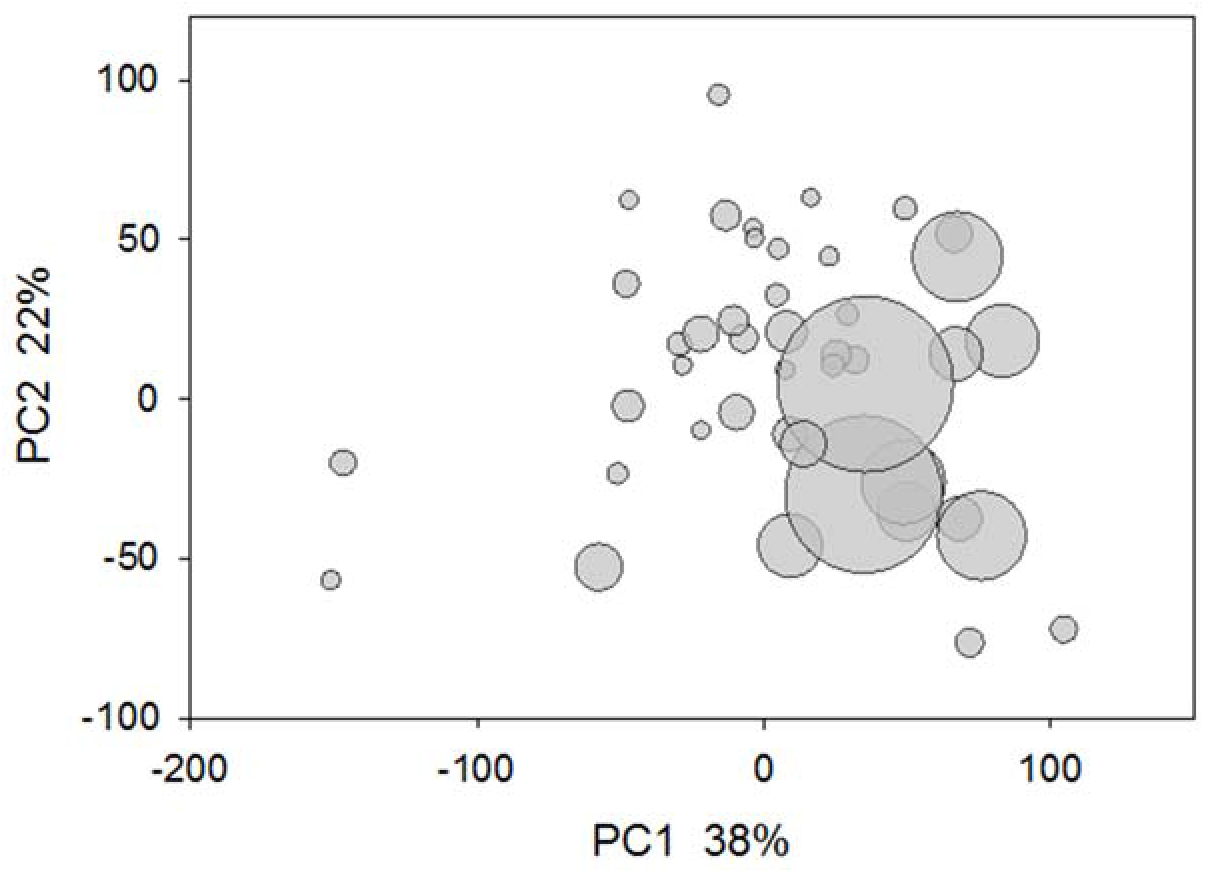
Principal Coordinates Analysis based on KEGG level 2 functional categories. Datapoints located in close proximity have similar metagenomes. The size of the circles is proportional to oocyst concentration. Metagenome gene abundance from experiment 2 was inferred from 16S sequence data using PICRUSt for 44 fecal samples analyzed by 16S amplicon sequencing and oocyst concentration.

**Table S1.** Metagenomic differences grouped by KEGG level 2 pathways between severely infected and mildly infected mice.

| Group | KEGG pathway | log(abundance) | LDA score | p |
| --- | --- | --- | --- | --- |
| Severely infected | Poorly Characterized | 4.704 | 2.80 | 0.034 |
|  | Cellular Processes and Signaling | 4.629 | 3.23 | 0.001 |
|  | Metabolism of Cofactors and Vitamins | 4.592 | 2.96 | 0.041 |
|  | Cell Motility | 4.540 | 3.59 | 0.001 |
|  | Genetic Information Processing | 4.466 | 2.92 | 0.010 |
|  | Metabolism unclassified | 4.439 | 2.71 | 0.002 |
|  | Folding Sorting and Degradation | 4.341 | 2.67 | 0.023 |
|  | Signal Transduction | 4.332 | 3.28 | 0.000 |
|  | Glycan Biosynthesis and Metabolism | 4.306 | 2.86 | 0.002 |
|  | Infectious Diseases | 3.656 | 2.61 | 0.001 |
|  | Circulatory System | 0.665 | 2.11 | 0.008 |
|  | Cardiovascular Diseases | 0.148 | 2.60 | 0.008 |
| Mildly infected | Carbohydrate Metabolism | 5.039 | 3.59 | 0.005 |
|  | Amino Acid Metabolism | 4.939 | 2.89 | 0.016 |
|  | Replication and Repair | 4.924 | 3.38 | 0.003 |
|  | Translation | 4.722 | 3.27 | 0.002 |
|  | Nucleotide Metabolism | 4.586 | 3.04 | 0.005 |
|  | Lipid Metabolism | 4.484 | 2.91 | 0.002 |
|  | Xenobiotics Biodegrad. & Metabolism | 4.336 | 3.05 | 0.003 |
|  | Metabolism of other Amino Acids | 4.200 | 2.53 | 0.028 |
|  | Cell Growth and Death | 3.685 | 2.38 | 0.001 |
|  | Signaling Molecules and Interaction | 3.353 | 2.25 | 0.001 |

